# Mathematical models for cell migration with real-time cell cycle dynamics

**DOI:** 10.1101/238303

**Authors:** Sean T. Vittadello, Scott W. McCue, Gency Gunasingh, Nikolas K. Haass, Matthew J. Simpson

## Abstract

Fluorescent ubiquitination-based cell cycle indicator, also known as FUCCI, allows the visualisation of the G1 and S/G2/M cell cycle phases of individual cells. FUCCI consists of two fluorescent probes, so that cells in the G1 phase fluoresce red and cells in the S/G2/M phase fluoresce green. FUCCI reveals real-time information about cell cycle dynamics of individual cells, and can be used to explore how the cell cycle relates to the location of individual cells, local cell density, and different cellular microenvironments. In particular, FUCCI is used in experimental studies examining cell migration, such as malignant invasion and wound healing. Here we present new mathematical models which can describe cell migration and cell cycle dynamics as indicated by FUCCI. The *fundamental* model describes the two cell cycle phases, G1 and S/G2/M, which FUCCI directly labels. The *extended* model includes a third phase, early S, which FUCCI indirectly labels. We present experimental data from scratch assays using FUCCI-transduced melanoma cells, and show that the predictions of spatial and temporal patterns of cell density in the experiments can be described by the fundamental model. We obtain numerical solutions of both the fundamental and extended models, which can take the form of travelling waves. These solutions are mathematically interesting because they are a combination of moving wavefronts and moving pulses. We derive and confirm a simple analytical expression for the minimum wave speed, as well as exploring how the wave speed depends on the spatial decay rate of the initial condition.

## 1 Introduction

The cell cycle consists of a sequence of four distinct phases, namely: gap 1 (G1); synthesis (S); gap 2 (G2); and the mitotic (M) phase [1]. The phases G1, S and G2 are collectively referred to as interphase, and involve cell growth and preparation for division. Following interphase, the cell enters the mitotic phase and divides into two daughter cells. While morphological changes associated with cell division can be observed visually during the transition from M to G1, such distinct morphological changes are not possible during transitions between other cell cycle phases [2]. Therefore, different techniques are required to study these other cell cycle transitions.

Since 2008, fluorescent ubiquitination-based cell cycle indicator (FUCCI) technology [2] has enabled the visualisation of the cell cycle progression from G1 to S/G2/M in individual cells. The FUCCI system consists of two fluorescent probes in the cell nucleus, or cytoplasm, which emit red fluorescence when the cell is in the G1 phase, or green fluorescence when the cell is in the S/G2/M phase. Prior to the development of FUCCI it was difficult, if not impossible, to examine the cell cycle dynamics of individual cells beyond the M to G1 transition [2]. In contrast, FUCCI allows direct visualisation, in real time, of transitions in the cell cycle. This technology is particularly useful for research in cancer biology [3, 4, 5, 6], cell biology [7, 8] and stem cell biology [9, 10].

Three-dimensional spheroids and two-dimensional scratch assays are commonly-used experimental models to study the invasive and proliferative behaviour of cancer cells. In combination with FUCCI, these experimental models can be used to examine the cell cycle dynamics of individual cells as a function of position within the spheroid or scratch assay [3, 5, 6]. A major advantage of this method is that two fundamental phenomena associated with malignant invasion, namely cell proliferation and cell migration, can be characterised simultaneously. Previous methods to examine the roles of cell migration and cell proliferation involve pre-treating cells with anti-mitotic drugs, such as mitomycin-C [11]. A major limita tion of these previous methods is that the application of the anti-mitotic drug is thought to suppress proliferation without interrupting migration. However, this assumption is questionable, and rarely examined [12]. The development of FUCCI technology obviates the need for such crude methods to isolate the roles of cell migration and cell proliferation. Instead, FUCCI allows us to directly examine the spatial and temporal patterns of cell proliferation within a migrating population. To the best of our knowledge, there are no mathematical models in the literature that have been developed to describe cell migration with FUCCI technology. The focus of this work is on cell *migration,* by which we mean a moving front of a population of cells. These moving fronts are composed of a large number of individual cells that do not maintain cell-to-cell contacts. The formation of the moving front of cells is driven by a combination of cell motility and cell proliferation.

Cell migration involves diffusion, arising from random cell motility, and proliferation of cells [12]. Mathematical models describing these processes in a population of cells tend to involve reaction-diffusion equations [13], which are often based on the Fisher–Kolmogorov–Petrovskii-Piskunov (FKPP) equation [14, 15],

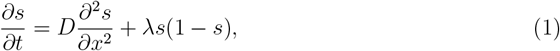

where *s*(*x*, *t*) > 0 is the cell density, *D* > 0 is the diffusion coefficient, and *λ* > 0 is the proliferation rate. Here, the dimensional cell density is scaled by the dimensional carrying capacity density, so that the maximum non-dimensional cell density is *s*(*x*, *t*) = 1. Carrying capacity limited proliferation of cells is described in Eq (1) with a logistic source term. Eq (1) has been successfully adapted to model many biological processes, such as *in vitro* cell migration [16, 17, 18]. A limitation of Eq (1) is that it considers a single population of cells. For a more realistic situation, where the total population is composed of a number of distinct, interacting subpopulations, it is relevant to consider a model that involves a system of coupled equations that are often related to Eq (1) [19].

The existence of travelling wave solutions for the FKPP equation is well known [15, 20]. Constant shape, monotonically decreasing wavefront travelling wave solutions propagate with a minimum speed, 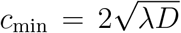 [15, 20]. Travelling wave solutions are of interest, because experimental observations tend to exhibit moving fronts that can be thought of as a travelling wave [16]. The speed of a moving cell front is often the simplest quantitative measurement that can be obtained from an experiment [16, 21]. Therefore, understanding the relationship between the parameters in the mathematical model and the speed of the travelling wave solution is a useful way to help parameterise the mathematical model to match experimental observations. Travelling wave solutions have also been observed in other mathematical models of cell migration [16, 19, 21, 22], as well as other reaction-diffusion models related to biological processes [23, 24, 25, 26]. Most travelling wave solutions take the form of moving wavefronts, which have a monotone profile. Another type of travelling wave solution is a pulse, which is characterised by a nonmonotone profile [23, 27].

Here we present a new mathematical model of cell migration, which incorporates cell cycle dynamics based on the information provided by FUCCI technology in relation to the cell cycle phase. We consider the cells in a particular phase of the cell cycle to make up a distinct subpopulation, so our model consists of a system of coupled partial differential equations. To motivate our generalisation of Eq (1), we pay careful attention to the underlying biological features. This leads us to develop two different mathematical descriptions of cell migration with FUCCI technology:

- **Fundamental FUCCI model:** In the most fundamental format, FUCCI highlights a subpopulation of cells in the G1 phase as being red, and another subpopulation of cells in the S/G2/M phase as being green. Motivated by the ability to distinguish between these two phases of the cell cycle, we develop a mathematical model with two subpopulations: *v_r_*(*x*, *t*) and *v_g_*(*x*, *t*). Here, the *v_r_*(*x*, *t*) subpopulation corresponds to the red cells and the *v_g_*(*x*, *t*) subpopulation corresponds to the green cells. We refer to this model as the *fundamental* model.
- **Extended FUCCI model:** In some experimental descriptions, cell biologists identify an additional subpopulation that corresponds to the situation where both of the red and green probes are active simultaneously, giving rise to a third subpopulation that appears to be yellow. This overlap of the red and green fluorescence occurs during the early S, or eS, phase. Using experimental images we find that the yellow subpopulation is more difficult to reliably identify than either the red or green subpopulations, as only a very small proportion of the cell population appear to be distinctly yellow. This is, in some sense, expected, because the yellow subpopulation results from the transient overlap of the G1 phase (red) and the S/G2/M phase (green). Despite this difficulty, we also develop another mathematical model which is capable of representing the three subpopulations: *u_r_*(*x*, *t*) is the red subpopulation, *u_y_(*x*, *t*)* is the yellow subpopulation, and *u_g_*(*x*, *t*) is the green subpopulation. We refer to this model as the *extended* model.

The fundamental and extended models are both related to the FKPP model with the dependent variable *s*(*x*, *t*). To summarise the similarities and differences between the fundamental and extended models, the FKPP model, and the relationship between these mathematical models and the underlying biology, we report the dependent variables and their biological interpretation in Table 1.

**Table 1.** Summary and comparison of the fundamental and extended models and the FKPP model.

| Model | Equation reference | Dependent variables | Biological interpretation |
| --- | --- | --- | --- |
| FKPP | Eq (1) | $s(x, t)$ | Total cell density is a function of position $x$ and time $t$ |
| Fundamental<br>FUCCI | Eqs (2)–(3) | $v_r(x, t)$ | Cell density for cells in G1 (red) phase is a function of position $x$ and time $t$ |
| | | $v_g(x, t)$ | Cell density for cells in S/G2/M (green) phase is a function of position $x$ and time $t$ |
| | | $s(x, t)$ | Total cell density, $s(x, t) = v_r(x, t) + v_g(x, t)$, is a function of position $x$ and time $t$ |
| Extended<br>FUCCI | Eqs (4)–(6) | $u_r(x, t)$ | Cell density for cells in G1 (red) phase is a function of position $x$ and time $t$ |
| | | $u_y(x, t)$ | Cell density for cells in eS (yellow) phase is a function of position $x$ and time $t$ |
| | | $u_g(x, t)$ | Cell density for cells in S/G2/M (green) phase is a function of position $x$ and time $t$ |
| | | $s(x, t)$ | Total cell density, $s(x, t) = u_r(x, t) + u_y(x, t) + u_g(x, t)$, is a function of position $x$ and time $t$ |

In this work, we present information relating to both the fundamental and the extended models, but we focus on developing new mathematical analysis of the fundamental FUCCI model. Furthermore, we quantitatively apply the fundamental model to new experimental data. There are two types of parameters in our model, namely transition rates between phases of the cell cycle, and diffusion coefficients describing the rate of cell migration. The transition rates between phases of the cell cycle are estimated using experimental data from Haass et al. [3], who report data relating to the time spent in each phase of the cell cycle. Using new experimental data in the form of images of scratch assays of FUCCI-transduced melanoma cells, we extract quantitative information about cell density as a function of position and time, and compare these quantitative data with the predictions of our fundamental model. Since this is the first time that a mathematical model has been used to predict a cell migration experiment with FUCCI-transduced cells, we focus on two-dimensional cell migration experiments as this is the most common experimental platform because of convenience, simplicity and low cost [28]. Two-dimensional cell migration assays are valuable because they are often used as high-throughput screening tools in conjunction with more sophisticated preclinical models [28]. In addition to showing how the new mathematical models can be used to predict the two-dimensional experiments, numerical solutions of the mathematical model show the formation of travelling wave solutions that are a combination of coupled wavefronts and pulses. We also derive an analytical expression for the minimum wave speed of the travelling waves, and we explore how these results are different from those of the standard FKPP equation, given in Eq (1).

This manuscript is organised as follows. In the Materials and Methods section we detail the experimental protocol used to perform the scratch assays using FUCCI-transduced melanoma cells. We also outline the numerical analysis of our mathematical models. In the Results and Discussion section we present and discuss our mathematical models, and compare numerical solutions of the fundamental model with new data from scratch assays of melanoma cells. We then analyse our models numerically for travelling wave solutions, and for the fundamental model we derive an analytical expression for the minimum wave speed.

## 2 Materials and methods

### 2.1 Experiments

**Cell culture**The human melanoma cell lines C8161 (kindly provided by Mary Hendrix, Chicago, IL, USA), 1205Lu and WM983C (both kindly provided by Meenhard Herlyn, Philadelphia, PA, USA) were genotypically characterized [29, 30, 31, 32], grown as described [33] (using 4% fetal bovine serum instead of 2%), and authenticated by STR fingerprinting (QIMR Berghofer Medical Research Institute, Herston, Australia).

#### Fluorescent ubiquitination-based cell cycle indicator (FUCCI)

To generate stable melanoma cell lines expressing the FUCCI constructs, mKO2-hCdt1 (30-120) and mAG-hGem (1-110) [2] were subcloned into a replication-defective, self-inactivating lentiviral expression vector system as previously described [33]. The lentivirus was produced by cotransfection of human embryonic kidney 293T cells. High-titer viral solutions for mKO2-hCdt1 (30/120) and mAG-hGem (1/110) were prepared and used for co-transduction into the melanoma cell lines, and subclones were generated by single cell sorting [3, 5, 34].

#### Wound healing migration assay

Experiments were performed using the three melanoma cell lines C8161, 1205Lu and WM983C. For each cell line, three independent experiments were performed. FUCCI-transduced melanoma cells from each cell line were seeded in a 6-well plate to subconfluence. The seeding density was adjusted according to the doubling time for the cell line. The monolayer was scraped with a p200 pipette tip, and images were taken at regular time intervals.

### 2.2 Mathematical model

#### Numerical solutions

Numerical solutions of Eqs (2)–(3) and Eqs (4)–(6) are obtained on a domain, 0 ≤ *x* ≤ *L*, with grid spacing Δ*x*, and with uniform time steps of duration Δ*t*.

The details are in Supporting Material 1.

In this study, the initial condition takes one of three forms, depending on the purpose of the modelling exercise:

- **Modelling a scratch assay:** The first set of modelling results involves using the fundamental mathematical model to mimic a set of experimental data from a scratch assay. For this purpose, we take images from the experiment, manually count numbers of cells in each phase of the cell cycle, and use these numbers to specify *v_r_*(*x*, 0) and *v_g_*(*x*, 0). To simulate the experiment, we solve the governing equation numerically on a finite domain, 0 ≤ *x* ≤ *L*, where *L* is chosen to match the physical dimension of the experimental image.
- **Exploring the minimum wave speed of travelling wave solutions:** The second set of modelling results involves studying long-time numerical solutions of the mathematical model to examine the possibility of travelling wave solutions. To ensure that we focus on the most biologically-relevant travelling-wave solutions, we apply initial conditions with compact support. Further details are provided in the Results and Discussion.
- **Dispersion relation:** Having demonstrated the existence of travelling wave solutions, we then analyse how the long-time travelling wave speed depends on the decay rate of the initial condition for the fundamental model. Further details are provided in the Results and Discussion.

## 3 Results and Discussion

### 3.1 Experimental data

Our experimental data come from two-dimensional scratch assays performed with FUCCI-transduced melanoma cells. In particular, we use the C8161, 1205Lu and WM983C melanoma cell lines [3]. Here we provide analysis for the C8161 and 1205Lu cell lines, which have very different cell cycle dynamics. We use these two cell lines to demonstrate that our model can predict the cell density for cell lines with a wide range of transition rates. The analysis for the WM983C cell line is in Fig S1 in Supporting Material 1. The WM983C cell line has cell cycle dynamics intermediate between the C8161 and 1205Lu cell lines.

In these experiments, melanoma cells migrate into a gap created by scratching the cell monolayer, and cell proliferation acts to increase the density of the monolayer. Still images of the scratch assays are obtained at four time points after the scratch is made: 0 h, 6 h, 12 h and 18 h for the C8161 cell line, see Fig 1A–D; 0 h, 16 h, 32 h and 48 h for the 1205Lu cell line, see Fig 2A–D; and 0 h, 16 h, 32 h and 48 h for the WM983C cell line, see Fig S1A–D. From these images, the nuclei of individual cells can be observed as red (G1 phase), yellow (eS phase) or green (S/G2/M phase). Over the time period of the experiments, the cells migrate into the gap, and cell proliferation is evident from the increasing density of cells behind the moving front. A notable feature of each of the images is that very few cells appear to be distinctly yellow. Almost all of the yellow cells appear to be either partly green and yellow, or partly red and yellow. This ambiguity motivates us to work with the fundamental mathematical model, which treats the yellow eS phase as part of the red G1 and green S/G2/M phases.

**Figure 1:**
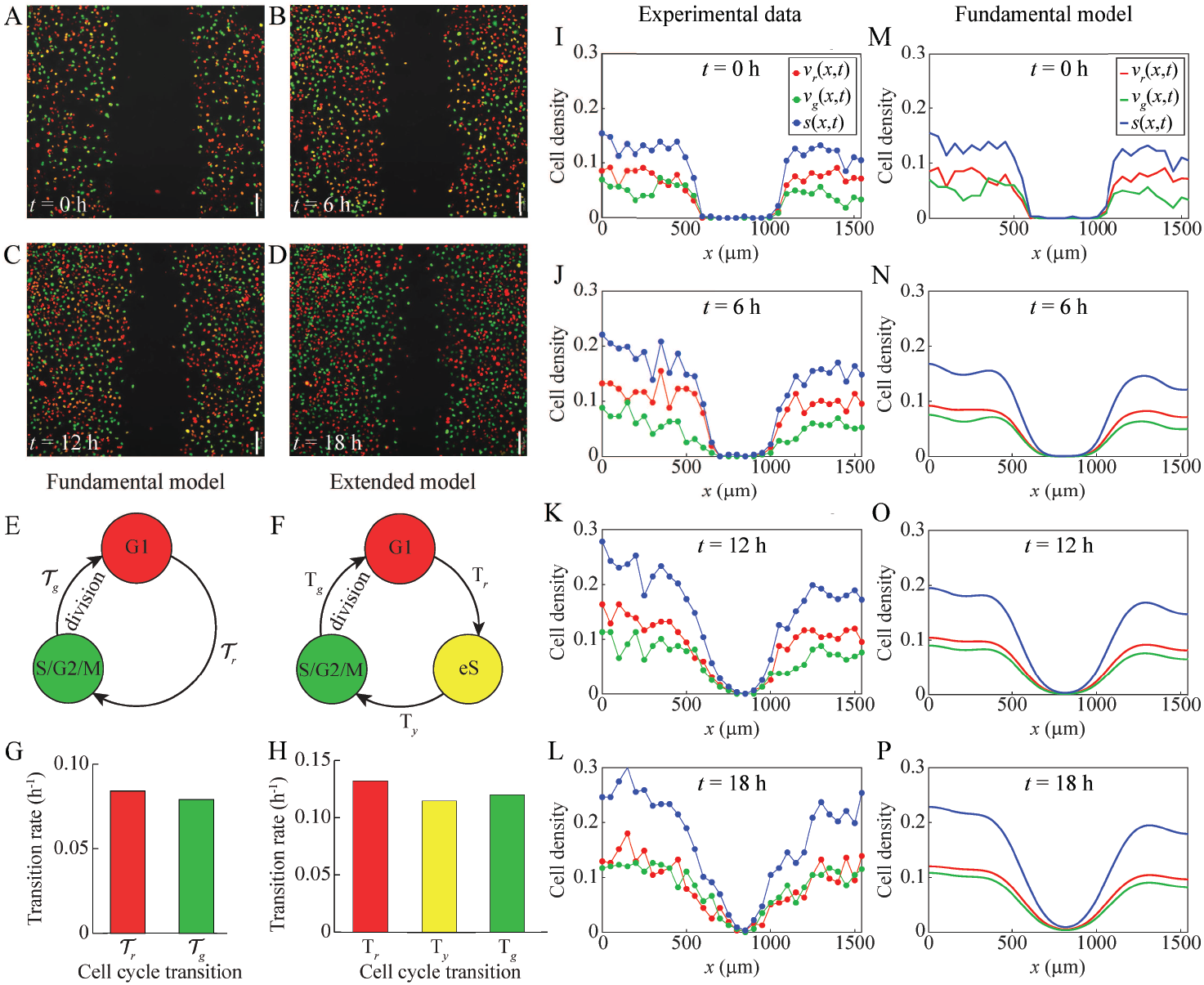
Comparison of experimental data and the fundamental model for a scratch assay of FUCCI-transduced C8161 melanoma cells. (A)–(D) Still images of a scratch assay with FUCCI-transduced C8161 melanoma cells at time 0 h, 6 h, 12 h and 18 h, respectively. Scale bar is 150 *μ*m. (E) Schematic of the fundamental model with two subpopulations indicating the transitions between the cell cycle phases indicated by FUCCI. (F) Schematic of the extended model with three subpopulations indicating the transitions between the cell cycle phases indicated by FUCCI. (G) Estimated transition rates from one cell cycle phase to the next for the fundamental model with two subpopulations, based on data from the C8161 cell line from Fig 1C in [3]. (H) Estimated transition rates from one cell cycle phase to the next for the extended model with three subpopulations, based on data from the C8161 cell line from Fig 1C in [3]. (I)–(L) Experimental non-dimensional cell density data at 0 h, 6 h, 12 h and 18 h, respectively (based on images (A)–(D)). The cell density is treated as a function of *x* and *t* only, owing to the fact that the initial density does not depend on the vertical coordinate, *y*. (M)–(P) Numerical solutions of the fundamental model, Eqs (2)–(3), at 0 h, 6 h, 12 h and 18 h. The numerical solutions are obtained with Δ*x* = 0.1 *μ*m, Δ*t* = 0.1 h, *L* = 1542 *μ*m, diffusion coefficients 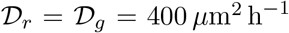, transition rates *κ_r_* = 0.084 h^−1^ and *κ_g_* = 0.079 h^−1^, and initial conditions the same as for the experimental data.

**Figure 2:**
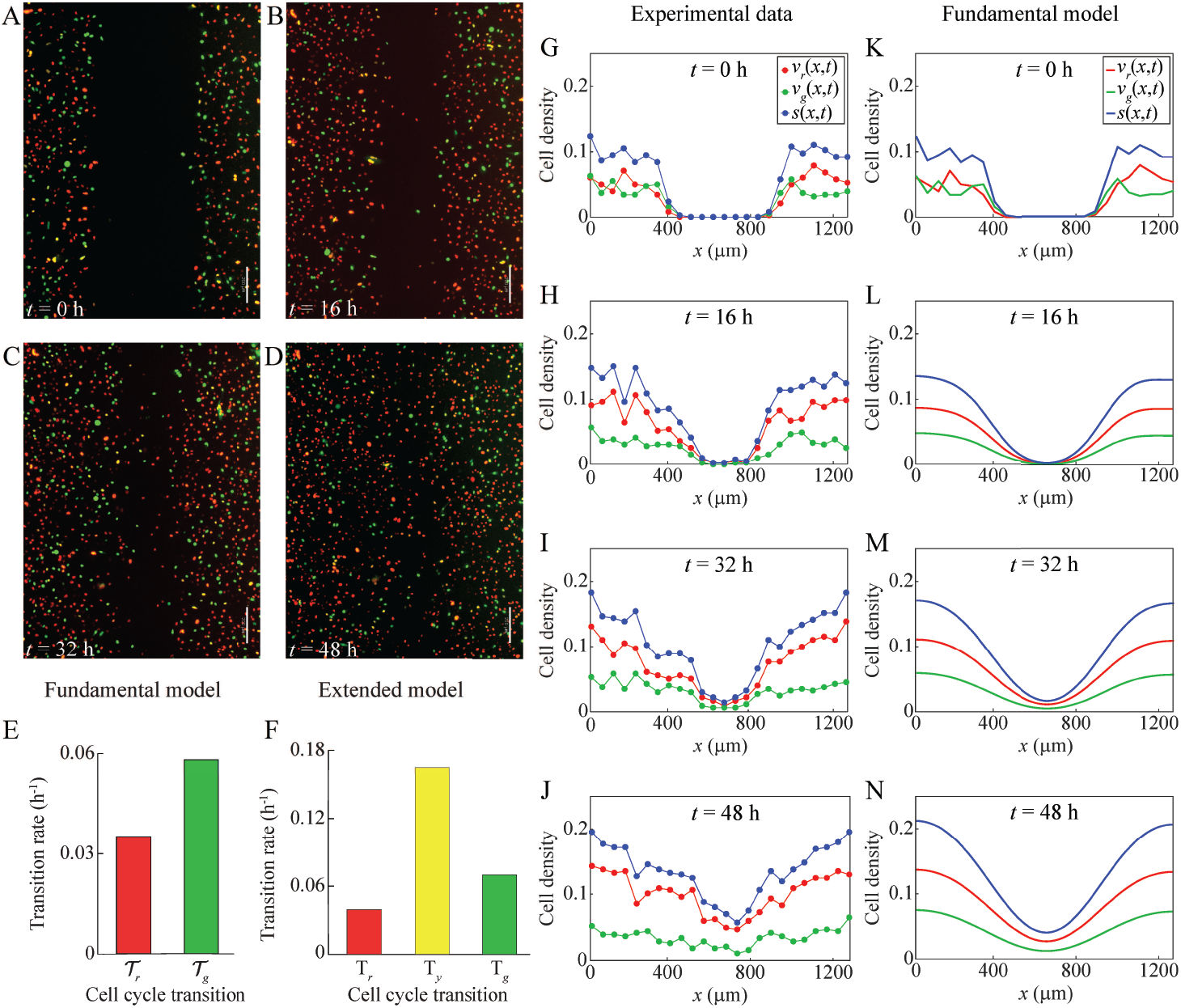
Comparison of experimental data and the fundamental model for a scratch assay of FUCCI-transduced 1205Lu melanoma cells. (A)–(D) Still images of a scratch assay with FUCCI-transduced 1205Lu melanoma cells at time 0 h, 16 h, 32 h and 48 h, respectively. Scale bar is 200 *μ*m. (E) Estimated transition rates from one cell cycle phase to the next for the fundamental model with two subpopulations, based on data from the 1205Lu cell line from Fig 1C in [3]. (F) Estimated transition rates from one cell cycle phase to the next for the extended model with three subpopulations, based on data from the 1205Lu cell line from Fig 1C in [3]. (G)–(J) Experimental non-dimensional cell density data at 0 h, 16 h, 32 h and 48 h, respectively (based on images (A)–(D)). The cell density is treated as a function of *x* and *t* only, owing to the fact that the initial density does not depend on the vertical coordinate, *y*. (K)–(N) Numerical solutions of the fundamental model, Eqs (2)–(3), at 0 h, 16 h, 32 h and 48 h. The numerical solutions are obtained with Δ*x* = 0.1 *μ*m, Δ*t* = 0.1 h, *L* = 1254 *μ*m, diffusion coefficients 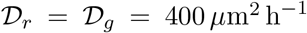, transition rates *κ_r_* = 0.035 h^−1^ and *κ_g_* = 0.058 h^−1^, and initial conditions the same as for the experimental data.

While the images in Figs 1A–D, 2A–D and S1A–D provide some quantitative data about these particular scratch assays for these particular cell lines, the purpose of using a mathematical model is to provide significant generalisations beyond what is possible when working purely with experiments. For example, if we use this kind of data to parameterise a mathematical model, then we ought to be able to use the parameterised mathematical model to make predictions about varying different aspects of the experiment, such as changing the width of the scratch, the timescale of the experiment, or the initial density of the monolayer. To parameterise our mathematical model we require cell density data from the experiment. While cells are free to move in two dimensions, the geometry of the experiment is such that the cell density is spatially uniform in the vertical, *y*, direction. We therefore quantify the cell density as a function of horizontal position, *x*, at various times, *t* [36]. Overall, this geometrical simplification allows us to describe the cell density data as a function of one spatial coordinate only, and we can therefore use a one-dimensional mathematical model to describe this kind of data [36].

To obtain cell density data from the images in Figs 1A–D, 2A–D and S1A–D, we divide each image into a series of vertical columns, each of width 50 *μ*m. We manually count the number of cells of each colour in each column. As previously discussed, there is some degree of ambiguity in classifying a cell as yellow, as almost all of the yellow cells appear to be a mixture of either red and yellow, or a mixture of green and yellow. This ambiguity is probably due to the fact that yellow arises from the transient overlap of red and green fluorescence. Consequently, we take the most straightforward approach and classify all of the cells as being either red or green. In this way we can work with just two subpopulations. Given the cell counts, we divide each cell count in each column by the area of that column to give the dimensional, column-averaged, cell density. These estimates of dimensional cell density are then converted into estimates of non-dimensional cell density by dividing through by the carrying capacity density, *K*. These data are provided in Supporting Material 2. To estimate *K* [37], we assume that the cells are uniformly sized disks, and that the maximum monolayer density corresponds to hexagonal packing of cells. Hexagonal close packing corresponds to one cell at each vertex of the hexagon, and one cell at the center of the hexagon, meaning that the hexagon contains the equivalent of three cells. The area of the hexagon is 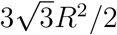, where *R* is the circumradius. If the radius of the cells is *a* then, since *R* = 2*a*, the carrying capacity is given by 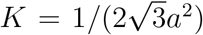. The cell diameter of C8161 melanoma cells is approximately 17 *μ*m [38]. So, with *a*= 8.5 *μ*m we have K = 0.004 cells *μ*m^−2^, to one significant figure. We use the same value of *K* for the 1205Lu and WM983C melanoma cells, as cells from the three cell lines are similar in size.

Fig 1I–L shows the resulting experimental cell density profiles for C8161 cells at 0 h, 6 h, 12 h and 18 h, respectively. Fig 2G–J shows the resulting experimental cell density profiles for 1205Lu cells at 0 h, 16 h, 32 h and 48 h, respectively. Similar profiles for the WM983C cells are in Fig S1G–J. These figures provide quantitative information about how the population of cells migrates into the gap, while simultaneously proliferating to increase the density of the spreading monolayer, as we observed qualitatively in Figs 1A–D, 2A–D and S1A-D. These cell density profiles, however, quantify the cell density of each subpopulation, and allow the changes in cell density over time to be determined quantitatively, which is not possible through visual interpretation of the images.

Movies of scratch assays associated with the three cell lines considered in this work are provided in Supporting Material 3–5.

### 3.2 Model development

We now describe the FUCCI scratch assays using the fundamental model. As previously explained, cells in the experiment move in two dimensions. However, the geometry of the experiment means that the cell density is spatially uniform in the vertical, *y*, direction. Therefore, we can model the experiment with a one-dimensional model, where the independent variables are time, *t*, and the horizontal coordinate, *x* [36].

As summarised in the Introduction, in the FUCCI system, red and green fluorescent proteins are fused to different regulators of the cell cycle so that a cell in G1 phase fluoresces red, and a cell in S/G2/M phase fluoresces green. During the cell cycle transition from G1 phase to S phase, referred to as eS phase, the red FUCCI signal decreases and the green FUCCI signal increases, producing varying shades of yellow fluorescence, ranging from darker yellow to lighter yellow. The indication of the eS (yellow) phase is therefore secondary, as it arises from the overlap of red and green fluorescence. In the experimental images, it is difficult to identify the cells in eS phase as very few cells appear distinctly yellow, rather appearing shades of either red or green. For these reasons, our fundamental mathematical model is

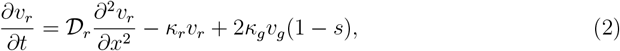

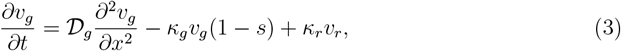

where *v_r_*(*x*, *t*) and *v_g_*(*x*, *t*) are the non-dimensional cell densities of the coupled subpopulations corresponding to the G1 (red) and S/G2/M (green) phases of the cell cycle, respectively. The total density is *s*(*x*, *t*) = *v_r_*(*x*, *t*) + *v_g_*(*x*, *t*). The rate at which cells in the G1 phase transition to the S/G2/M phase is *κ_r_*, and the rate at which cells in the S/G2/M phase transition to the G1 phase is *κ_g_*. The diffusion coefficients are 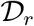 for cells in phase G1, and 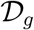 for cells in phase S/G2/M. While we are free to choose any realistic values for the diffusion coefficients, it is pertinent to set 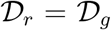, since we are considering subpopulations of cells of the same type which differ only with respect to their cell cycle phase. Indeed, cells from various melanoma cell lines, including C8161, 1205Lu and WM983C, appear to migrate independently of the cell cycle phases. For example, see the data in Figure 6 and Figure S3 in [3]. We will employ this biologically-motivated simplifying assumption at various points in our analysis. Note that in Eq (2), the factor of two in the positive source term corresponds to a cell in phase S/G2/M undergoing division to produce two daughter cells in the phase G1, thereby doubling the local density.

Despite the challenges associated with the observation of the yellow eS phase, we also consider a mathematical model for three coupled subpopulations corresponding to G1, eS and S/G2/M. We refer to this model as our extended model:

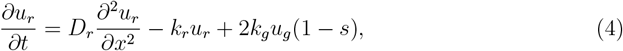

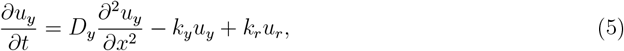

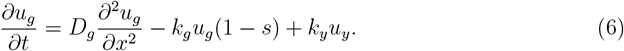

Here, *u_r_*(*x*, *t*), *u_y_*(*x*, *t*) and *u_g_*(*x*, *t*) are the non-dimensional cell densities of the coupled subpopulations corresponding to the G1 (red), eS (yellow) and S/G2/M (green) phases of the cell cycle, respectively, with total cell density *s*(*x*, *t*) = *u_r_*(*x*, *t*) + *u_y_*(*x*, *t*) + *u_g_*(*x*, *t*). The G1 to eS transition rate is *k*_*r*_, the eS to S/G2/M transition rate is *k_y_* and the S/G2/M to G1 transition rate is *k*_*g*_. The diffusion coefficients are *D_r_* for cells in phase G1, *D_y_* for cells in phase eS, and *D_g_* for cells in phase S/G2/M. Once again we have no restriction on the choice of the values for the diffusion coefficients. The subpopulations consist of cells of the same type, they are only in different phases of the cell cycle. So, as discussed above for the fundamental model, the most obvious choice is to set *D_r_* = *D_y_* = *D_g_*. As in Eq (2), the factor of two in Eq (4) corresponds to a cell in phase S/G2/M undergoing division to produce two daughter cells in the phase G1.

### 3.3 Model application

We illustrate our mathematical model in Figs 1, 2 and S1, where we compare experimental data for scratch assays of FUCCI-transduced melanoma cells with the fundamental model Eqs (2)–(3). Fig 1 corresponds to the C8161 melanoma cell line, Fig 2 to the 1205Lu melanoma cell line, and Fig S1 to the WM983C melanoma cell line.

Fig 1E describes, schematically, the fundamental model with two subpopulations, indicating the transitions between the cell cycle phases indicated by FUCCI. We denote the transitions between the cell cycle phases as

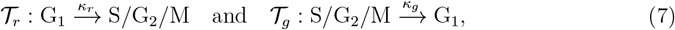

where *κ_r_* is the G1 to S/G2/M transition rate, and *κ_g_* is the S/G2/M to G1 transition rate. Fig 1F is a similar schematic for the extended model with three subpopulations, where we denote the transitions between the cell cycle phases as

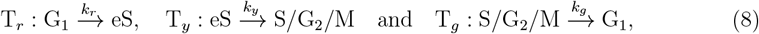

where *k_r_*, *k_y_* and *k_g_* are the G1 to eS, eS to S/G2/M and S/G2/M to G1 transition rates, respectively.

The estimated transition rates from one cell cycle phase to the next phase are based on FUCCI data for the C8161, 1205Lu and WM983C melanoma cell lines from Fig 1C in [3]. These data report the duration of time spent in each cell cycle phase for at least 20 individual cells. To estimate the transition rate from one cell cycle phase to the next, we first calculate the arithmetic mean of the data in [3], giving the mean times: *t_r_* for the G1 phase, *t_y_* for the eS phase, and *t_g_* for the S/G2/M phase. We then estimate the transition rates for the extended model as *k_r_* = (ln 2)/*t_r_* for the G1 to eS transition, *k_y_* = (ln 2)/*t_y_* for the eS to S/G2/M transition, and *k_g_*= (ln 2)/*t_g_* for the S/G2/M to G1 transition. The factor of ln2 arises since this data corresponds to cells in a low density environment, and so the cells are likely to be proliferating exponentially. To obtain estimates of the transition rates for the fundamental model, we assume that half of the time spent in the eS phase contributes to the time spent in the G1 phase, and the other half of the time spent in the eS phase contributes to the time spent in the S/G2/M phase. This means that we have *κ_r_* = (ln 2)/(*t_r_* + *t_y_*/2) for the G1 to S/G2/M transition, and *κ_g_* = (ln 2)/(*t_g_* + *t_y_*/2) for the S/G2/M to G1 transition. The estimated transition rates for the fundamental and extended models are shown in Figs 1G–H, respectively, for the C8161 cell line, in Figs 2E–F, respectively, for the 1205Lu cell line, and in Figs S1E–F, respectively, for the WM983C cell line. We can express the transition rates for the fundamental model in terms of those for the extended model as *κ_r_* = 2*k_r_k_y_*/(*k_r_* + 2*k_y_*) and *κ_g_* = 2*k_g_k_y_*/(*k_g_* + 2*k_y_*).

Previous studies examining the migration of various melanoma cell lines suggest that the diffusion coefficients lie within the range 100-500 *μ*m^2^h^−1^ [39, 40]. Therefore, we will take an intermediate value and assume that the diffusivity of our melanoma cell lines is approximately 400 *μ*m^2^h^−1^.

For comparison with the experimental data for C8161 cells in Fig 1I–L, we show the numerical solutions of the fundamental model, Eqs (2)–(3), in Fig 1M-P. Overall, the numerical solutions of Eqs (2)–(3) in Fig 1M-P compare well with the corresponding experimental data in Fig 1I-L. As time increases, the solution of the mathematical model shows the cell density profile spreading into the gap, and the cell density of each subpopulation increases throughout the domain due to proliferation. The relative size of each subpopulation also compares well between the numerical solutions and experimental data. Similarly, for comparison with the experimental data for 1205Lu cells in Fig 2G–J, we show the numerical solutions of the fundamental model, Eqs (2)–(3), in Fig 2K–N. Once again, the numerical solutions of Eqs (2)–(3) in Fig 2K–N compare well with the corresponding experimental data in Fig 2G–J. A similar comparison for the WM983C cell line is in Fig S1.

We note that other studies use partial differential equations to model scratch assays [16, 19, 36]. None of these previous models, however, include the cell cycle phases of cells. Our results show potential for our model to successfully describe cell migration and proliferation along with cell cycle dynamics. With our model we can easily simulate experiments which would otherwise be expensive and time consuming. In particular, we can simulate experiments over longer periods of time, with different scratch widths, and with different parameters to accommodate different cell lines. Our model also provides quantitative data such as cell densities at any time and position within the domain.

### 3.4 Analysis

#### Numerical solutions

A key feature that can be observed in scratch assays initialised with a sufficiently wide scratch is the formation of a moving front of cells [16]. This is important, because similar observations are relevant to malignant invasion and wound healing [16]. The need to understand the key factors that drive moving fronts of cells, motivates an analysis of our models for scenarios in which a single front propagates along a wide domain.

We now simulate solutions of our models on a much wider domain using a different initial condition to examine the existence of travelling wave solutions. There are many choices of initial condition; however, for the fundamental model we set

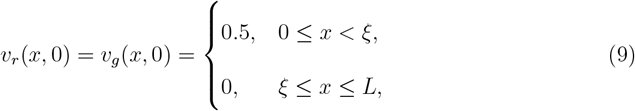

and for the extended model we set

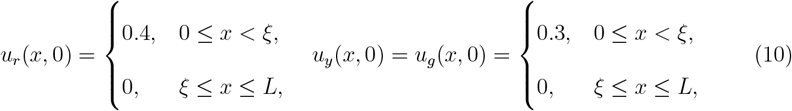

where the choice of the constant *ξ* is not critical. Since we use a numerical approach to explore travelling wave solutions, we set *L* to be sufficiently large so that the moving front does not interact with the boundary at *x* = *L*.

For this purpose, we obtain numerical solutions of the fundamental model, Eqs (2)–(3), with typical solutions presented in Fig 3A. These results suggest that the solutions develop into an interesting travelling wave profile, where the density of cells in the G1 phase forms a moving pulse, while the density of cells in the S/G2/M phase forms a moving wavefront. Furthermore, the total cell density profile also moves as a wavefront profile. Analogous solutions of the extended model, Eqs (4)–(6), are shown in Fig 3B, which have similar features except that there is an additional pulse arising from cells in the eS phase. We find that the existence of these travelling waves is robust, and does not depend on the values of the diffusion coefficients. Fig S2 shows solutions of the fundamental model for 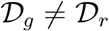.

**Figure 3:**
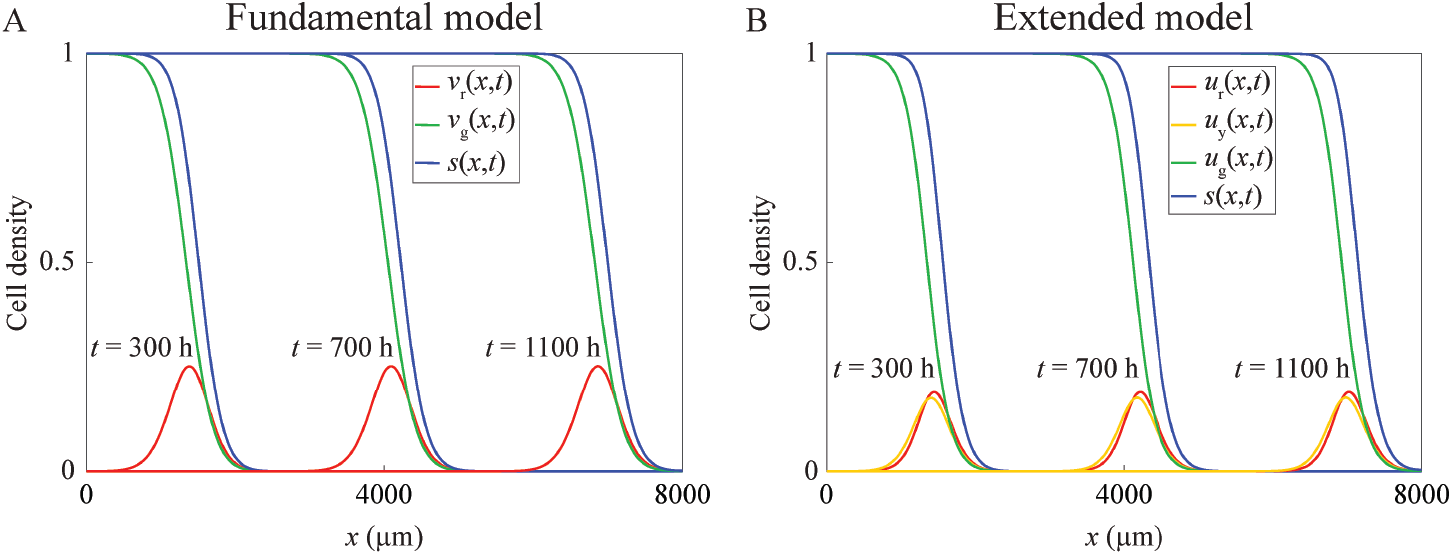
Numerical solutions demonstrating travelling wave behaviour for the fundamental and extended models. (A) Numerical solutions of the fundamental model, Eqs (2)–(3), obtained with Δ*x* = 1 *μ*m, Δ*t* = 1h, *L* = 8000 *μ*m, 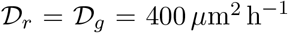, *κ_r_* = *κ_g_* = 0.08 h^−1^, and the initial condition given by Eq (9) with= 10. (B) Numerical solutions of the extended model, Eqs (4)–(6), obtained with Δ*x* = 1 *μ*m, Δ*t* = 1h, *L* = 8000 *μ*m, **D_r_**=**D_y_**=*D_g_*= 400 *μ*m^2^ h^−1^, *k_r_* = *k_y_* = *k_g_* = 0.13 h^−1^, and the initial condition given by Eq (10) with *ξ* = 10.

The appearance of travelling wave solutions which take the form of a wavefront is not unexpected, as the partial differential equations in our model are related to the FKPP equation, Eq (1), which is well-known to exhibit travelling-wave solutions with a wavefront form [15, 20]. It is particularly interesting, however, that our model also exhibits travelling wave solutions with the form of a pulse, which are not observed for the FKPP equation. The pulses arise because only the cells near to the leading edge, where *s*(*x*, *t*) < 1, have the opportunity to cycle from S/G2/M to G1, which involves cell division and can be inhibited by crowding effects. Behind the wavefront where *s*(*x*, *t*) approaches unity, cells do not have enough space to divide, and so these cells remain in the S/G2/M phase.

#### Travelling wave analysis of the fundamental model

We now analyse the fundamental model, Eqs (2)–(3), with the aim of understanding how the parameters in the model relate to the speed of the travelling wave solutions. To simplify our analysis we non-dimensionalise the problem by defining the new variables *t*\* = *κ_g_t* and 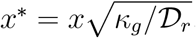, and the parameters to give the corresponding non-dimensional model

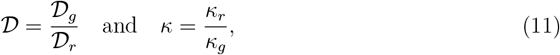

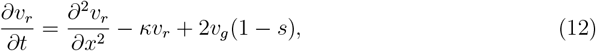

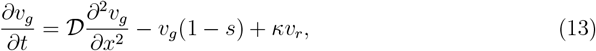

in which the asterisks have been omitted for notational simplicity.

To examine the travelling wave solutions, we define the travelling wave coordinate, *z* = *x* − *ct,* where *c* > 0 is the wave speed associated with a travelling wave that propagates in the positive x-direction. We seek solutions of Eqs (12)–(13) of the form *v_r_*(*x*, *t*) = *U*(*z*) and *v_g_*(*x*, *t*) = *V*(*z*). Such solutions, if they exist, correspond to travelling wave solutions. Substituting *U*(*z*) and *V*(*z*) into Eqs (12)–(13) gives the system of ordinary differential equations

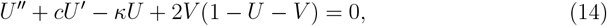

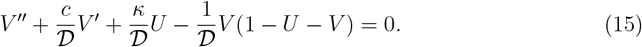

We want to find travelling-wave solutions which satisfy the conditions

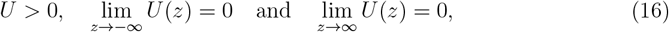

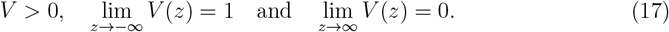

Letting *W* = *U′* and *X* = *V*′, we can write Eqs (14)–(15) as a system of first-order equations:

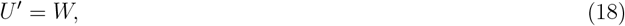

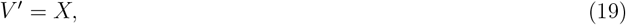

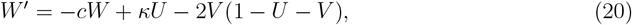

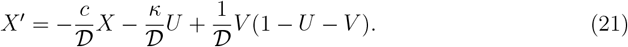

The equilibrium points of Eqs (18)–(21) are (0, 0, 0, 0) and (0,1, 0, 0). Of all the solutions to Eqs (18)–(21) in the four-dimensional phase space, (*U*, *V*, *W*, *X*), we seek a heteroclinic orbit from (0,1, 0,0) to (0, 0, 0, 0) which has the physically-relevant property that *U* > 0 and *V* > 0.

The Jacobian of Eqs (18)–(21) evaluated at (0, 0, 0, 0) is

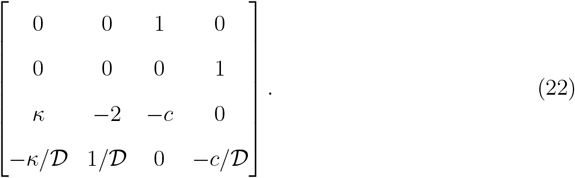

The eigenvalues of Eq (22) are the solutions of the corresponding characteristic equation

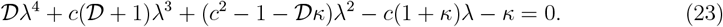

To establish a condition for physically-relevant travelling-wave solutions, where *U* > 0 and *V* > 0, we examine whether the solutions of Eq (23) are either complex or real. The analytical solutions of this quartic equation [41] are quite complicated when 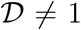. Since we are considering subpopulations of the same cell type which differ only with respect to cell cycle phase, the biologically-relevant case is when the two diffusion coefficients are equal, leading to 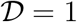. In this case, by defining

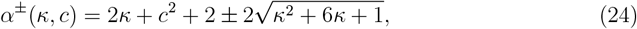

we can express the solutions of the quartic simply as

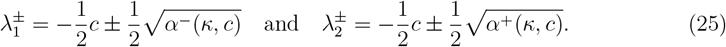

The eigenvalues 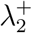 and 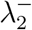 are always real, and 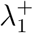 and 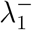 are real when *α*^−^(*κ*, *c*) ≥ 0. If *α*^−^(*κ*, *c*) < 0, however, then 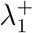 and 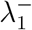 are complex eigenvalues which yield solutions with oscillatory behaviour, which necessarily involves negative cell densities. Since we are interested in travelling wave solutions for which *U* and *V* remain positive, we shall therefore require that *α*^−^(*κ*, *c*) ≥ 0. It is then reasonable to suspect that, for a given *κ* ≥ 0, the value of c such that *α*^−^(*κ*, *c*) = 0 is the minimum wave speed for the travelling waves, which we denote as

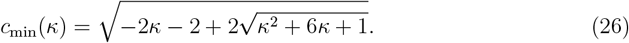

The minimum wave speed is bounded above, and in fact *c*_min_(*κ*) → 2^−^ as *κ* → ∞.

Eq (26) shows that the minimum speed of the travelling wave solution depends on *κ*, which is the ratio of the time the cells spend in phase S/G2/M to the time the cells spend in phase G1. In other words, the minimum wave speed depends on the cell cycle dynamics of the particular cells under consideration. We observe here that *α*^−^(*κ*, *c*) ≥ 0 is a necessary condition for the existence of travelling waves. We have not demonstrated, however, that this condition is sufficient for the existence of travelling waves. This would require a formal proof of existence for the travelling waves, which is beyond the scope of this current work.

Observe that if *κ* > 0 and *c* ≥ *c*_min_ then 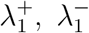 and 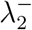 are real and negative, and 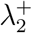 is real and positive. Therefore, the equilibrium point (0, 0, 0, 0) is hyperbolic and has a three-dimensional stable manifold and a one-dimensional unstable manifold. The presence of a stable manifold at the point (0, 0, 0, 0) is necessary for the existence of a heteroclinic orbit and the real eigenvalues allow for this orbit to correspond to physically relevant travelling wave solutions with *U* > 0, *V* > 0. This analysis does not constitute a formal proof of existence for the travelling waves, however it does show that our observations are consistent with their existence.

The Jacobian of the system Eqs (18)–(21) evaluated at (0, 1, 0, 0) is

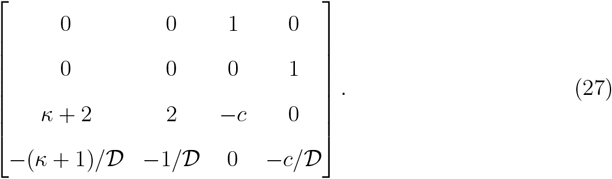

The eigenvalues of Eq (27) are the solutions of the corresponding characteristic equation

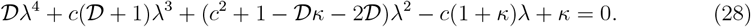

Once again, the analytical solutions of this quartic equation [41] are quite complicated when 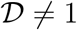. For the biologically-relevant case 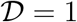, however, the solutions are

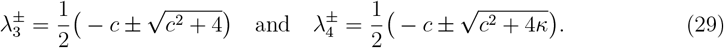

If *κ* > 0 then 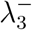 and 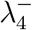 are real and negative, and 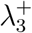 and 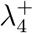 are real and positive. Therefore, the equilibrium point (0,1, 0, 0) is hyperbolic and has a two-dimensional stable manifold and a two-dimensional unstable manifold. The existence of an unstable manifold at the point is necessary for the presence of a travelling wave solution.

Fig 4A compares *c*_min_(*κ*) in Eq (26) with the wave speed estimates obtained from the numerical solutions of the partial differential equations [21]. The numerical solutions are obtained using initial conditions with compact support, so we would expect the resulting travelling waves to have the minimum wave speed [15]. We observe that the numerically-estimated wave speed is consistent with Eq (26) over the range of *κ* we consider. Therefore, we have numerical evidence to strongly support the claim that the minimum speed is given by Eq (26).

**Figure 4:**
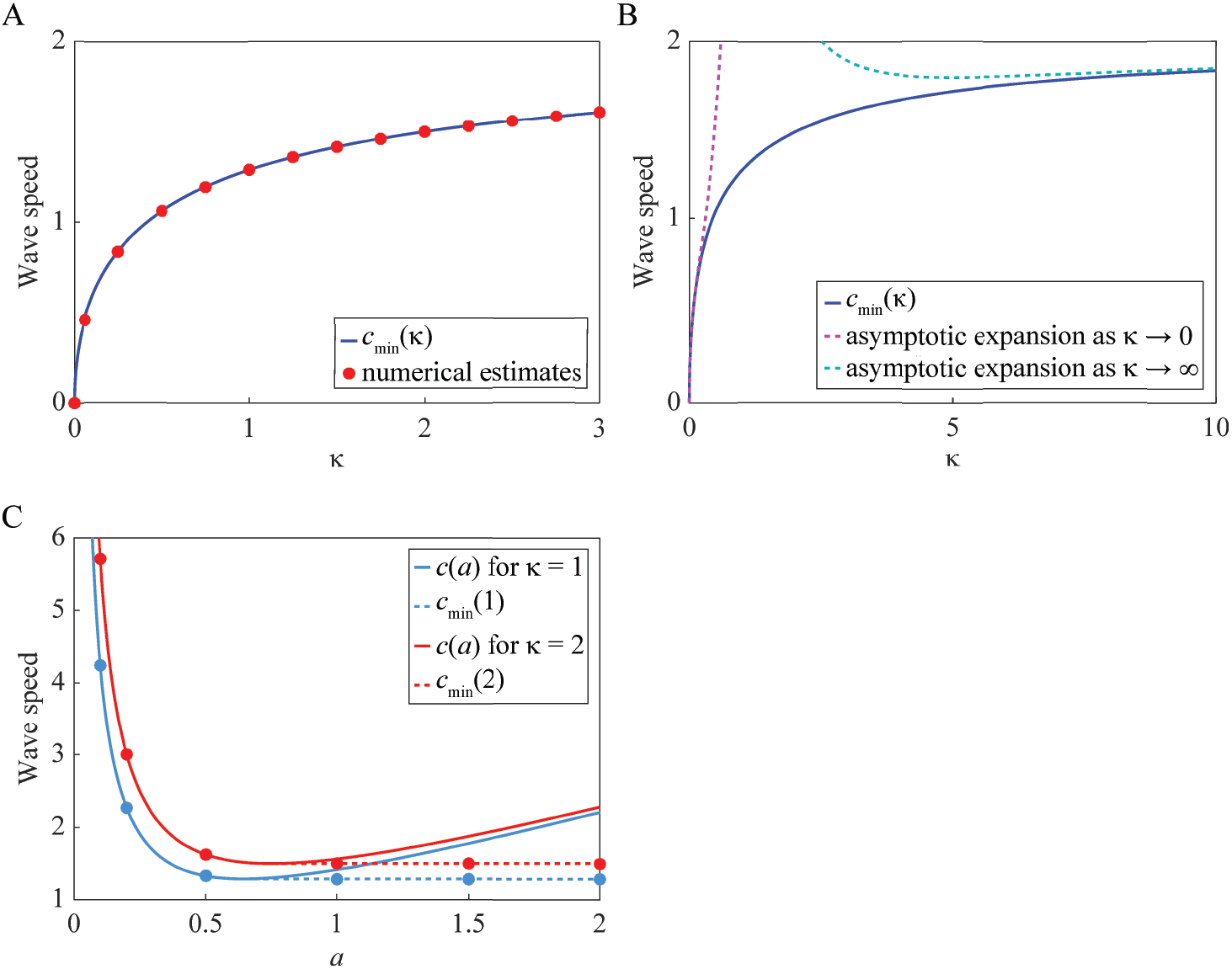
Minimum wave speed, *c*_min_(*κ*), and the dispersion relation. (A) Comparison of *c*_min_(*κ*) from Eq (26) with the numerically-estimated wave speed for *s*(*x*, *t*) = *v_r_*(*x*, *t*) + *v_g_*(*x*, *t*). Numerical solutions of Eqs (12)–(13) are obtained with Δ*x* = 0.1, Δ*t* = 0.001 and 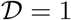. Further, the initial condition is Eq (9) with *ξ* = 10. For *κ* = 0 there is no travelling wave, so we set *c* = 0. The numerical solutions are considered beginning with *κ* = 0.06, and then with increasing values of *κ* from 0.25 to 3, with increments of 0.25. From these numerical solutions we estimate the wave speed for s(*x*, *t*) = *v_r_*(*x*, *t*) + *v_g_*(*x*, *t*) by using linear interpolation to find the position corresponding to *s*(*x*, *t*) = 0.5 on the wave for various times [21]. (B) Asymptotic expansions for *c*_min_(*κ*) as *κ* → 0 and *κ* → ∞. (C) Comparison of *c* from Eq (35) with the wavespeed estimated using numerical solutions and with *c*_min_ from Eq (26). Solutions are given for *κ* = 1 (blue) and *κ* = 2 (red). The continuous curves show c from Eq (35). The dots represent the wave speed from numerical solutions obtained with Δ*x* = 0.1, Δ*t* = 0.001, 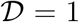, and initial conditions of the form Eq (36) with *α*= 0.1, 0.2, 0.5, 1, 1.5 and 2. The dotted horizontal lines show *c*_min_ from Eq (26).

In Fig 4B we show the asymptotic expansions of Eq (26) as *κ* → 0 and *κ* → ∞:

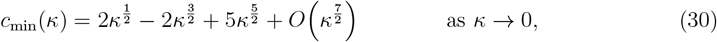

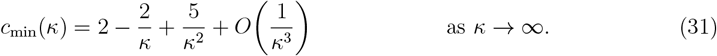

Thus, *c*_min_(*κ*) behaves like 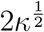 when kis small and like 2 − 2/*κ* when *κ* is large. We discuss the connection with the FKPP equation in the Conclusion.

#### Dispersion relation

Here we investigate the relationship between the initial conditions for Eqs (12)–(13) with 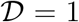, and the speed of the resulting travelling wave solution. Our approach is to examine the leading edge of the travelling wave, assuming that the initial conditions at infinity have an exponential form [15].

At the leading edge of the evolving waves, *s*(*x*, *t*) = *v_r_*(*x*, *t*) + *v_g_*(*x*, *t*) ≪ 1, so we can linearise Eqs (12)–(13) to give

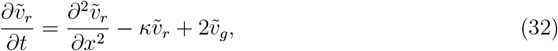

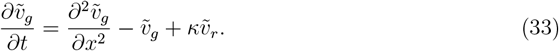

Assuming initial conditions of the form 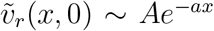 and 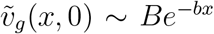 as *x* → ∞ for arbitrary positive constants *a*, *b*, *A* and *B*, we seek travelling wave solutions satisfying Eqs (32)–(33) of the form

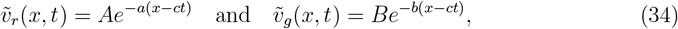

corresponding to the leading edges of the pulse and wavefront solutions, respectively. Substituting Eq (34) into Eqs (32)–(33) and solving for *c* gives

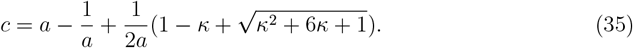

The dispersion relation, Eq (35), depends only on *a* and *κ*, so we can obtain the travelling wave solutions with the form Eq (34) from initial conditions of the form

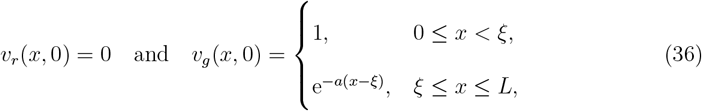

for constants *ξ* and *a* > 0. Note that for large *a*, this initial condition for *v_g_* is approximately a Heaviside function. For a given *κ*, we observe that the minimum wave speed according to the dispersion relation, Eq (35), is equal to the minimum wave speed *c*_min_, given by Eq (26).

In Fig 4C we compare c from Eq (35) with estimates of the wave speed from numerical solutions of the governing partial differential equations, and with *c*_min_ from Eq (26). The value of c given by the dispersion relation tends to infinity as both *a* → 0 and *a* → ∞, and has a unique minimum value for a given *κ*. The dispersion relation for the FKPP equation has similar properties [15]. For a given *κ*, the numerical estimates of the wave speed agree with Eq (35) for increasing values of *a* > 0, until the minimum wave speed is obtained. As we further increase *a*, the numerical estimates of the wave speed remain constant, at the minimum value of the wave speed. Once again, the dispersion relation for the FKPP equation has similar properties [15].

## 4 Conclusion

Here we present the first mathematical model of cell migration that can be used to quantitatively describe experiments using FUCCI technology, which highlights the spatial and temporal distribution of individual cells in different parts of the cell cycle. The fundamental model consists of two coupled partial differential equations, each of which governs the subpopulations of cells corresponding to the two phases of the cell cycle that are directly labelled by FUCCI. Our study suggests that the model can describe cell migration and cell proliferation in a way that highlights the spatial and temporal distribution of two subpopulations. In particular, we show that the model can describe the dynamics of scratch assays performed with cells highlighted by FUCCI. This is a useful outcome, as scratch assays are routinely employed to study cell migration, for example in the context of malignant invasion [36] and wound healing [16]. While a typical scratch assay may require several days to grow cells, and to perform and record the experiment, our model can simulate such an experiment on a single desktop computer in a few seconds. In addition, we can easily vary the parameters in our model to simulate experiments over any period of time, any scratch width, any geometry, and any cell line, provided that we have some information about the amount of time that is spent in each phase of the cell cycle. Therefore, this kind of computational modelling tool can provide valuable information to assist in the design and interpretation of these kinds of experiments conducted with FUCCI.

In this work we use numerical results to demonstrate the existence of travelling wave solutions. Furthermore, our analysis shows that the minimum wave speed depends on the ratio of the time spent in each of the G1 and S/G2/M phases. Another outcome of this study is that we derive an analytical expression for the minimum wave speed as a function of this ratio which, in dimensional variables, can be written as

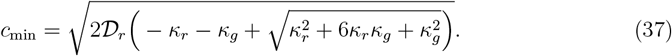

This relationship is based on the biologically-reasonable assumption that 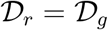, where 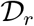 is the diffusion coefficient for cells in phase G1 and 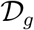 is the diffusion coefficient for cells in phase S/G2/M. Further, *κ_r_* is the transition rate from G1 to S/G2/M and *κ_g_* is the transition rate from S/G2/M to G1. It then follows from Eqs (30)–(31) that 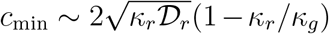 as *κ_r_*/*κ_g_* → 0, and 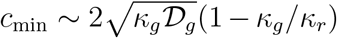 as *κ_g_*/*κ_r_* → 0. Therefore, when *κ_r_*/*κ_g_* ≪ 1, so that cells spend much more time in phase G1 compared with phase S/G2/M, the minimum wave speed obtained from our fundamental FUCCI model, Eqs (2)–(3), approaches the minimum wave speed obtained from the FKPP equation, Eq (1). A similar observation holds for the case *κ_g_*/*κ_r_* ≪ 1, corresponding to the situation where the cells spend much more time in phase S/G2/M compared with phase G1.

Travelling wave solutions are of great practical interest as cell migration tends to exhibit travelling wave characteristics [16]. The analytical expression we derive for the minimum wave speed is of practical interest, as a moving front of cells can be thought of as acting like a travelling wave solution, so our expression can provide a prediction for the speed of the moving front in experimental studies. The travelling wave solutions of the fundamental model are mathematically interesting because they are a combination of moving wavefronts and moving pulses. Monotonically decreasing wavefront solutions are well-known for the FKPP equation, and since our models are related to the FKPP equation, it is not surprising that wavefront solutions are also observed in our study. It is interesting, however, that travelling wave solutions of our models involve pulses, which are not features of the travelling wave solutions of the FKPP equation.

There are many possibilities for future work arising from this study. An area of particular interest would involve using our models to quantitatively study how the migration of melanoma cells and the cell cycle for melanoma cells are affected by the application of particular melanoma drugs. Indeed, there is still much to learn regarding the effects of introduced drugs on melanoma cell activity [3]. These kinds of drugs often act to arrest the cell cycle, thereby preventing melanoma proliferation. Another feature that could be examined is the role of contact inhibition and cell cycle arrest, since it is accepted that cells in relatively high density environments can undergo cell cycle arrest. Indeed, our model does not account for this phenomenon, since we are interested in low to moderate cell densities. Another way that this study could be extended is to consider additional phases of the cell cycle. This is of interest because a very recent extension of FUCCI technology, referred to as FUCCI4 [42], highlights all four cell cycle phases G1, S, G2 and M. If extended to four coupled partial differential equations, our modelling framework could be used to quantitatively describe cell migration where individual cells are highlighted using FUCCI4.

## 5 Acknowledgments

We thank Emeritus Professor Sean McElwain and two anonymous referees for helpful comments. NKH is a Cameron fellow of the Melanoma and Skin Cancer Research Institute, Australia, and is supported by the National Health and Medical Research Council (AP1084893). MJS is an Australian Research Council Future Fellow, and is supported by the Australian Research Council Discovery Program (DP170100474). SWM is supported by the Australian Research Council Discovery Program (DP140100249).

